# Structure and immune recognition of the porcine epidemic diarrhea virus spike protein

**DOI:** 10.1101/2020.02.18.955195

**Authors:** Robert N. Kirchdoerfer, Mahesh Bhandari, Olnita Martini, Leigh M. Sewall, Sandhya Bangaru, Kyoung-Jin Yoon, Andrew B. Ward

## Abstract

Porcine epidemic diarrhea virus is an alphacoronavirus responsible for significant morbidity and mortality in pigs. A key determinant of viral tropism and entry, the PEDV spike protein is a key target for the host antibody response and a good candidate for a protein-based vaccine immunogen. We used electron microscopy to evaluate the PEDV spike structure, as well as pig polyclonal antibody responses to viral infection. The structure of the PEDV spike reveals a configuration similar to that of HuCoV-NL63. Several PEDV protein-protein interfaces are mediated by non-protein components including a glycan at Asn264 and two bound palmitoleic acid molecules. The polyclonal antibody response to PEDV infection shows a dominance of epitopes in the S1 region. This structural and immune characterization provides new insights into coronavirus spike stability determinants and explores the immune landscape of viral spike proteins.

## Introduction

Porcine epidemic diarrhea virus is a coronavirus of the alphacoronavirus genus. Identified as a viral agent distinct from transmissible gastroenteritis virus (Wood, 1977) and as a coronavirus (Pensaert and de Bouck, 1978), this virus is responsible for an enteric infection in pigs. Originally identified in England (Wood, 1977), PEDV is now a global pathogen. PEDV was first identified in the United States in 2013 (Stevenson et al., 2013) where it swept through pig populations causing several million piglet deaths (Sawyer and Sherwell, 2014). Mortality due to viral infection varies with age, with mortalities of 1-3 day old piglets in naïve herds approaching 100% (Lee, 2015). While mortality due to PEDV infection is lower in older pigs, infection can still result in decreased growth performance.

Like all coronaviruses, PEDV possesses a spike glycoprotein (S) responsible for cell attachment and virushost membrane fusion mediating viral entry into host cells. Coronavirus spikes are class I viral fusion proteins (Bosch et al., 2003; Chambers et al., 1990), possessing structural and functional parallels to influenza hemagglutinin (Wilson et al., 1981) and HIV-1 Env (Julien et al., 2013; Lyumkis et al., 2013). This class of proteins proceeds from a metastable prefusion conformation to a highly stable postfusion conformation (Bullough et al., 1994). For coronavirus spikes, this transition of conformations has been proposed to be triggered by progressive destabilization of the prefusion structure through receptor binding and host proteolytic cleavage (Belouzard et al., 2009; Millet and Whittaker, 2015; Taguchi and Matsuyama, 2002). As the major surface glycoprotein on the enveloped virions, spikes are also the target of neutralizing antibodies (Okda et al., 2017). These antibodies primarily target the PEDV S1 region and some epitopes have been found to be neutralizing (Li et al., 2017a; Okda et al., 2017; Sun et al., 2007). However, a structural mapping of PEDV antibody epitopes targeted during viral infection is lacking.

Coronavirus spikes adopt a shared domain arrangement where the N-terminal S1 regions of the homotrimeric spike surround and cap the C-terminal S2 regions. In betacoronavirus spikes, the S1 and S2 regions are often demarcated by a protease cleavage site (Millet and Whittaker, 2015). However, this site is lacking in many alphacoronvirus spikes including PEDV. Coronavirus S1 regions of the spike contain domains contributing to receptor binding. Receptors include both host glycans and proteins and vary widely between coronaviruses. PEDV has been shown to bind to sialic acid glycans using its S1 domain 0 (Li et al., 2016). By analogy with TGEV, PEDV spikes were also proposed to recognize porcine aminopeptidase N (pAPN) (Li et al., 2007). However, the use of pAPN as a receptor for PEDV has been called into question (Li et al., 2017b).

High-resolution structural studies of coronavirus spike proteins have the potential to shed light on molecular mechanisms of spike prefusion stability and the lessons learned can be utilized in the production of stabilized spike proteins as vaccine immunogens. Here we used cryoelectron microscopy (cryo-EM) to determine the structure of the PEDV spike protein at 3.5 Å resolution. This structure has allowed us to compare our findings with the structure of the human alphacoronavirus NL63 spike as well as another recently published PEDV spike structure (Walls et al., 2016b; Wrapp and McLellan, 2019). We found several protein-protein interfaces mediated by non-protein components suggesting roles for these glycans and ligands in enhanced prefusion spike stability. We also used negative-stain electron microscopy to illuminate the recognition of the PEDV spikes by the host polyclonal antibody response and map dominant antibody epitopes on the spike surface. Together, these data supply insights for the production of prefusion stabilized spikes as vaccine immunogens.

## Results

### Structural description

Coronavirus spikes are homotrimeric glycoproteins which can be separated into an N-terminal S1 region containing the receptor binding regions and a C-terminal S2 region containing the membrane fusion machinery (Fig. 1A). The overall structure of the PEDV spike strongly resembles that of the human coronavirus NL63 spike (Walls et al., 2017) (Fig 1B-D, Supplementary Table 1 and Supplementary Fig. 1). Like the NL63 spike, PEDV spike contains two structurally homologous S1 N-terminal domains, domain 0 and domain A. The arrangement of these S1 N-terminal domains is similar to that of NL63 where domain A occupies a position at the apex of the spike, distal to the viral membrane, while domain 0 is positioned beneath domain A, packing against the side of the spike protein and contacting both S1 domain D (also called sub-domain 2) and S2. The observed PEDV domain arrangement is in contrast to the recently published PEDV spike structure where domain 0 has flipped up and away from the main portion of the spike (Wrapp and McLellan, 2019) (Supplementary Fig. 2). Though sequence differences do exist between this previous PEDV spike structure (strain CV777) and that presented here (strain USA/Colorado/2013), these differences are small compared to the sequence differences with HuCoV-NL63 which is structurally more consistent with the PEDV spike described here. Indeed, even in our attempts at 3D classification of the PEDV spike particles, we do not see evidence for this flipped up conformation of domain 0. To test the previously proposed hypothesis that the difference in domain 0 conformations may be due to differences in expression system, we expressed the PEDV spike in both insect and mammalian cells including expression conditions to produce high-mannose glycosylation patterns similar to the methods used for the previous PEDV spike structure (Wrapp and McLellan, 2019). In all cases, we observe domain 0 in the beneath-domain A conformation (Supplementary Table 2 and Supplementary Fig. 2). The reasons for the altered conformation observed in the previous PEDV spike structure remain to be determined and will require additional experimental studies of alphacoronavirus spike structure.

**Figure 1:**
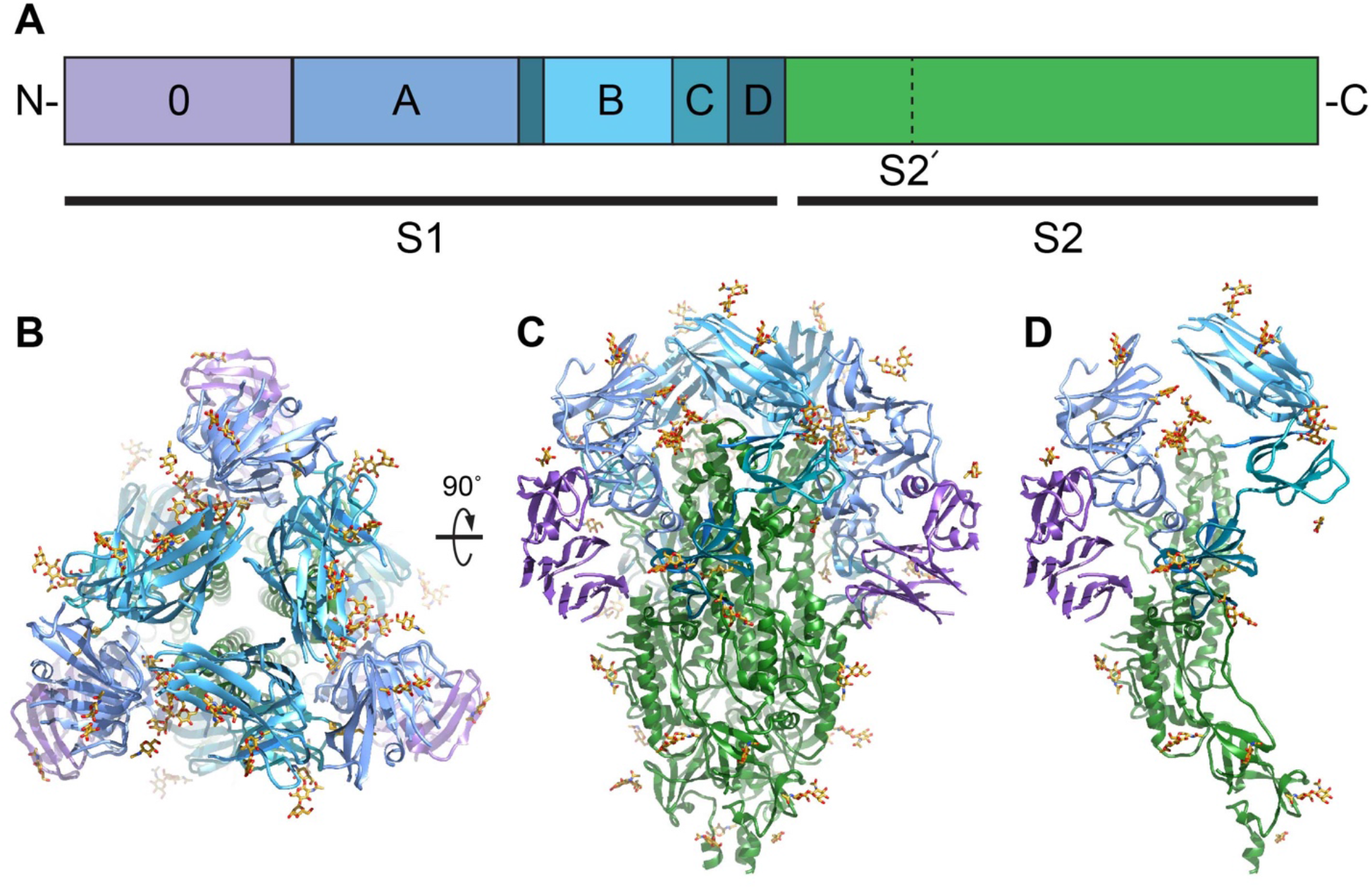
Overall structure of the PEDV spike protein. A) The primary sequence of the PEDV spike can be divided into S1 receptor-binding and S2 fusion machinery regions based on homology with betacoronaviruses. The S1 region can be further subdivided into domains 0, A, B, C and D (purple, light blue, cyan, teal and dark teal respectively. The S2 fusion machinery contains the S2’ cleavage site N-terminal to the fusion peptide. B) Viewing the trimeric PEDV spike protein from the membrane distal apex shows that all three copies of the domain B are in a downwards conformation. C) A 90° view of the spike trimer and D) and the corresponding view of the spike monomer demonstrate that PEDV adopts a conformation highly similar to other coronavirus spikes particularly HuCoV-NL63 (Walls et al., 2016b).

**Figure 2:**
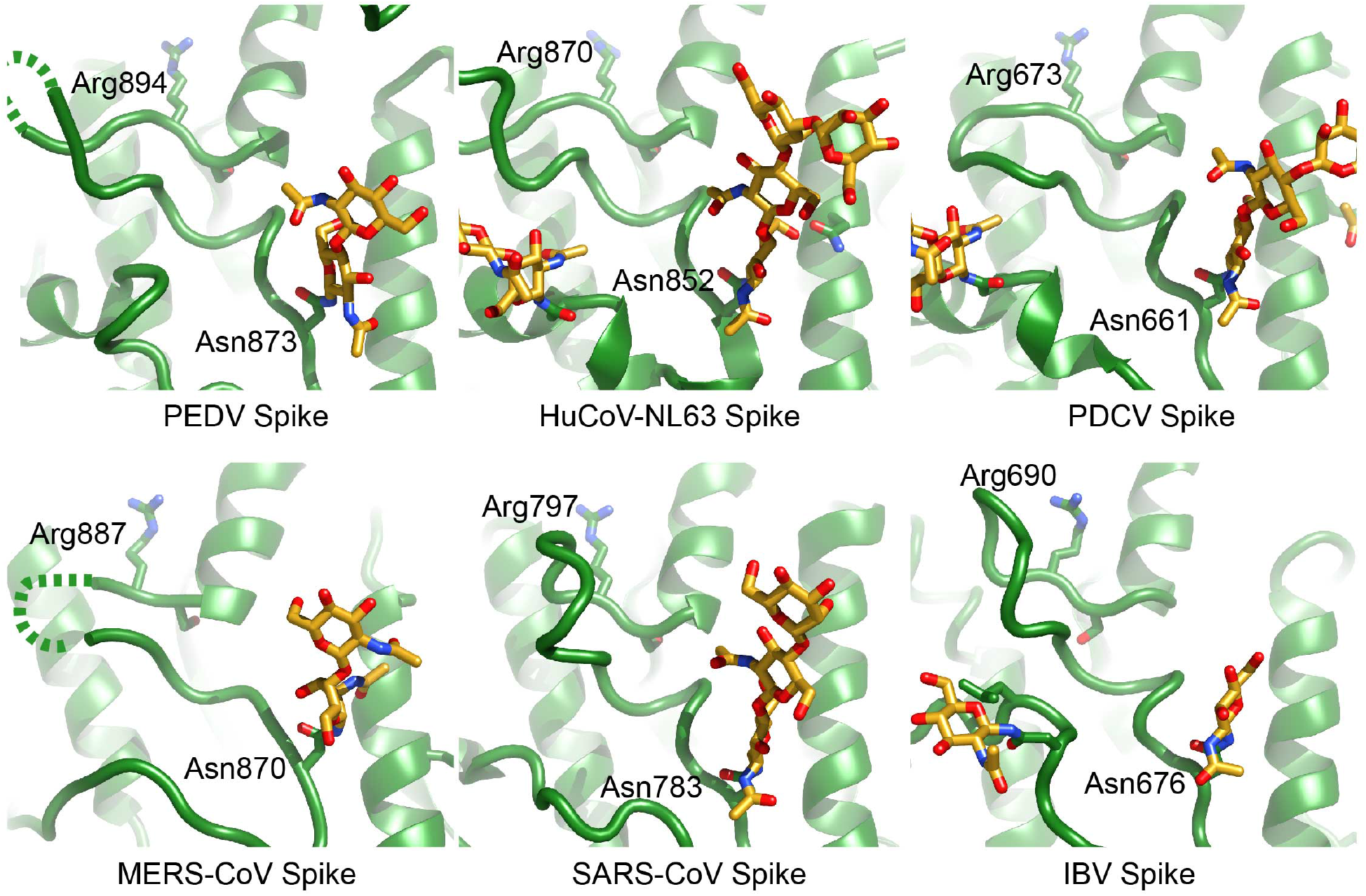
Comparison of S2’ cleavage sites in coronavirus spike structures. S2’cleavage occurs after a coronavirus conserved arginine residue. This position is preceded by protein loop and an N-linked glycan which appear to occlude this cleavage site in the prefusion conformation. Structures used for comparison include spikes from HuCoV-NL63 (5SZS.pdb (Walls et al., 2016b)), Porcine deltacoronavirus (6BFU.pdb, (Xiong et al., 2018)), MERS-CoV (5W9I.pdb, (Pallesen et al., 2017)), SARS-CoV (6CRV.pdb, (Kirchdoerfer et al., 2018)) and Infectious bronchitis virus (6CV0.pdb, (Shang et al., 2018a)).

Despite superimposing well onto the S2 fusion subunit and containing a similar overall S1 domain arrangement, the PEDV spike differs in several regards to the previously determined NL63 spike structure. Unlike the NL63 spike, the reconstructed density for domain 0 is more poorly defined than other domains indicating a higher degree of motion relative to the rest of the spike. Superimposition of the coordinate models of NL63 spike and the PEDV spike presented here reveals that both domain 0 and domain A have shifted approximately 15 Å towards the S2 region in the PEDV spike while domain B (also referred to as the S1 CTD) has remained in approximately the same position. The three B domains of the trimer are also more tightly packed around the spike three-fold axis with domain B making fewer contacts to domain A than observed in NL63 spike. All three B domains are in the downwards or lying conformation similar to that observed for available alpha-, gamma-, and deltacoronavirus spike structures (Shang et al., 2018a; Shang et al., 2018b; Walls et al., 2016b) as well as several betacoronavirus spike structures (Kirchdoerfer et al., 2016; Tortorici et al., 2019; Walls et al., 2016a).

PEDV spike is known to bind sialic acid and hemagglutinate red blood cells. This activity has been attributed to the S1 domain 0 (Hou et al., 2017; Li et al., 2016). However, comparison of the PEDV S1 domain 0 with the equivalent domain of the recently determined betacoronavirus OC43 spike in complex with sialic acid (Tortorici et al., 2019) suggests that the OC43 spike sialic acid binding site is not shared with PEDV despite structural homology between the two glycan-binding domains indicating an altered mode of glycan recognition in PEDV. In a portion of field isolates and tissue culture adapted viruses, PEDV S1 domain 0 has been deleted, leading to an apparent loss of sialic acid binding activity (Diep et al., 2017; Hou et al., 2017; Masuda et al., 2015). Though originally suggested to result in viruses with reduced mortality in piglets (Hou et al., 2017; Masuda et al., 2015), spike domain 0 deletion viruses have been found to be capable of causing high mortality (Diep et al., 2017). Moreover, it has been suggested that the deletion of domain 0 may contribute to viral persistence and reoccurrence of PEDV on pig farms (Diep et al., 2017). To examine the effect of this domain deletion on the overall structure of the PEDV spike, we used negative-stain EM to determine the structure of the PEDV spike with a 197 amino acid deletion, removing domain 0 (Diep et al., 2017). This low-resolution structure displays a similar structure to the full-length spike indicating that the deletion of this domain does not impart any macroscopic changes in spike protein conformation (Supplement Fig. 1).

The S2’ protease cleavage site in the S2 region of the spike is conserved across coronavirus spikes (Millet and Whittaker, 2015) and is believed to only be exposed for cleavage by host proteases during viral entry (Belouzard et al., 2009; Park et al., 2016). Examination of the PEDV S2 fusion machinery indicates a high structural homology with other coronavirus spike proteins including the S2’ cleavage site. This S2’ cleavage site at Arg894 is presented on the side of the spike following a short loop. The conformation of this preceding loop along with an upstream glycan at Asn873, blocks recognition of the S2’ cleavage site by host proteases in the prefusion conformation (Fig. 2). A comparison of available coronavirus spike structures (Kirchdoerfer et al., 2018; Pallesen et al., 2017; Shang et al., 2018a; Tortorici et al., 2019; Walls et al., 2016b; Xiong et al., 2018) shows the positioning and conformation of the preceding loop (875-893) to be conserved. In addition, the presence of a glycan 18-22 amino acids N-terminal to the S2’ cleavage site also appears to be shared across coronavirus genera.

### Sugar and fatty acids mediate protein chain interfaces

In modeling coordinates into the reconstructed EM density, we observed three non-protein densities at different domain interfaces. The first of these is a glycan density emanating from Asn264. This glycan is also present in the NL63 spike structure previously determined (Walls et al., 2016b) and occupies a hole observed in coronavirus spikes. However, due to the lower positioning of the S1 domain A relative to S2, this glycan in PEDV spike is sandwiched between the S1 domain A and the S2 region of an adjacent protomer. In coronavirus spikes, the S1 domain B caps the S2 central helices. So far unique to PEDV is the additional coordination of these S2 central helices by the Asn264 glycan (Fig. 3A) suggesting a contributing role for this glycan in stabilizing the prefusion spike conformation. Indeed, the sequon coding for this glycan position is 99.8% conserved across full-length PEDV spike sequences in the NCBI protein database (Pickett et al., 2012). As our PEDV spike protein used for high-resolution cryo-EM was produced in insect cells which possess alternative glycosylation patterns (Thirstrup et al., 1993), we sampled the overall conformations of PEDV spikes possessing a variety of glycosylation patterns by expressing an identical PEDV spike DNA construct using mammalian HEK-293F, with kifunensine, and HEK-293S Gnt^-/-^ expression systems and examining the protein with negative-stain EM (Supplementary Fig. 1). The low-resolution reconstructions reveal identical conformations of the PEDV spike despite the differing glycosylation states, including the lower positioning of the S1 domain A relative to the S2 region. We also removed the sequon for the Asn264 glycan by mutating the Asn to Asp. This mutation strongly reduced PEDV spike protein expression levels in HEK-293F cells by approximately 10-fold, consistent with a stabilizing effect. Nonetheless, this spike protein was still capable of adopting the pre-fusion conformation (Supplementary Fig. 2) indicating that the presence of the Asn264 glycan is not strictly required.

**Figure 3:**
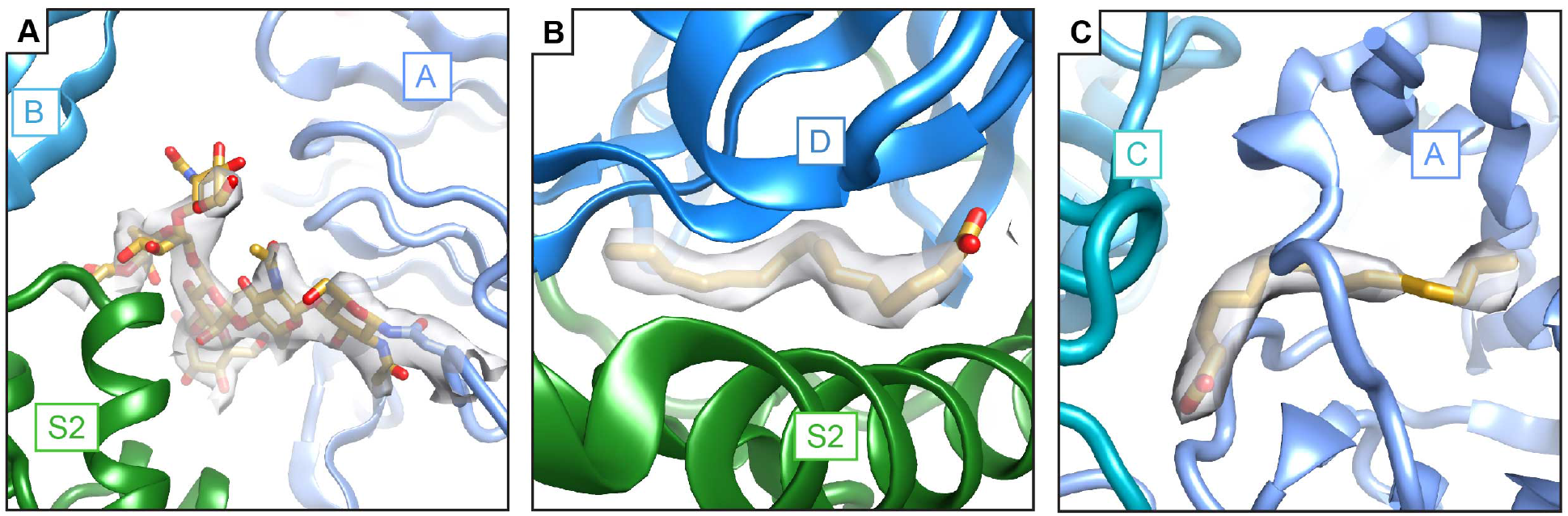
Non-protein components mediate protein-protein interactions in the PEDV spike. A) An N-linked glycan at Asn264 is sandwiched between domain A (light blue) and the S2 region of an adjacent protomer (green), capping the S2 central helices. B) Density for palmitoleic acid binding between S1 domain D (blue) and the S2 subunit of an adjacent protomer (green). B) A second site for palmitoleic acid was located in S1 domain A where the fatty acid head group reaches out to interact with S1 domain C of an adjacent protomer (teal).

The second non-protein density that we observed at an inter-domain interaction site lies between S1 domain D (also called sub-domain 2) and the S2 region of an adjacent protomer (Fig. 3B). The elongated, unbranched appearance of this density and a close proximity to neighboring Arg842 in S2 led us to model this density as a 16-carbon, unbranched fatty acid. An identical unmodeled density is present in the previously determined PEDV spike structure (Wrapp and McLellan, 2019). We confirmed the presence of this ligand using mass spectrometry, which indicated the presence of a molecule with a molecular weight matching that of palmitoleic acid (Supplementary Fig. 3). This fatty acid occupies a hydrophobic pocket formed between domain D and S2 and has limited surface exposure leading us to hypothesize that this fatty acid was incorporated during the folding of the protein domains. Comparison with published structures of NL63, 229E and Feline infectious peritonitis spikes shows that despite some amino acid sequence differences, the hydrophobic fatty acid pocket is shared between these alphacoronavirus spikes (Li et al., 2019; Walls et al., 2016b; Yuan et al., 2017). Moreover, examination of the NL63 spike reconstructed density (Walls et al., 2016b) reveals unmodeled density of a nearly identical ligand at this position (Supplementary Fig. 3) suggesting that the binding of palmitoleic acid to this site may be shared across alphacoronaviruses. It should be noted that while the reconstructed densities of 229E and feline infectious peritonitis virus spikes do contain some unmodeled density in this region, these smaller densities are less indicative of a stoichiometrically bound ligand. This binding pocket for palmitoleic acid is so far a feature only for available alphacoronavirus spike structures as spike structures from beta-, gamma- and deltacoronavirus spikes possess neither an intact hydrophobic pocket between domain D and S2 nor density for a bound ligand nearby. While the shared palmitoleic acid binding site between NL63 and PEDV spikes suggests an important role for this ligand, the importance of this binding site is enigmatic and remains to be explored experimentally. One possibility may be the burial of additional hydrophobic surfaces to stabilize interactions between S1 and S2.

A third non-protein density was observed in S1 domain A near its contact with the S1 domain C from an adjacent protomer (Fig. 3C). This density was also modeled as palmitoleic acid due to its positioning within a hydrophobic pocket and its unbranched appearance. A similar unmodeled density is present in the previously determined PEDV spike, albeit in altered conformation possibly owing to the different conformation of domains 0 and A in this structure (Wrapp and McLellan, 2019). In contrast to the palmitoleic acid binding pocket described above, this hydrophobic pocket is composed entirely of S1 domain A. However, the palmitoleic acid head group faces the S1 domain C from an adjacent protomer. Neighboring basic residues on the surface of domain C may provide favorable electrostatic interactions for the palmitoleic acid head group. Also, in contrast to the first palmitoleic acid site described above, density for this second palmitoleic acid is not present in the previously determined NL63 spike structure and is so far unique to the PEDV spike.

### Reactivity of PEDV spike with polyclonal antibody sera

To structurally characterize the antibody responses to PEDV infection, we examined serum samples from infected pigs for the presence of antibodies targeting PEDV spike proteins. Fab derived from the IgG of pigs experimentally infected with PEDV or a negative control animal was combined with recombinant PEDV spike and single particle negative-stain electron microscopy was performed as recently described (Bianchi et al., 2018). Extensive 3D classification revealed that all three of the sera from experimentally infected pigs had spike-specific antibodies that targeted the same two epitopes in S1 (Fig. 4, Supplementary Table 3 and Supplementary Fig. 4). The first epitope is located at the spike apex primarily contacting the membrane distal loops of S1 domain A as well as possibly contacting select loops of domain B. The second epitope lies on the side of the PEDV spike bridging S1 domains C and D. This side epitope of PEDV spike corresponds to the previously mapped “S1 D” neutralizing epitope, amino acids 636-789 (Okda et al., 2017; Sun et al., 2007). Antibodies recognizing this epitope were shown to contain much of the neutralizing activity of anti-PEDV polyclonal sera. Interestingly, we did not observe Fabs binding to other previously identified epitopes including domain 0 or domain B (Chang et al., 2002; Li et al., 2017a) possibly owing to a lower prevalence of these antibodies or the unique truncated immunogens used to originally elicit antibodies against these epitopes rather than intact trimeric spikes presented during infection.

**Figure 4:**
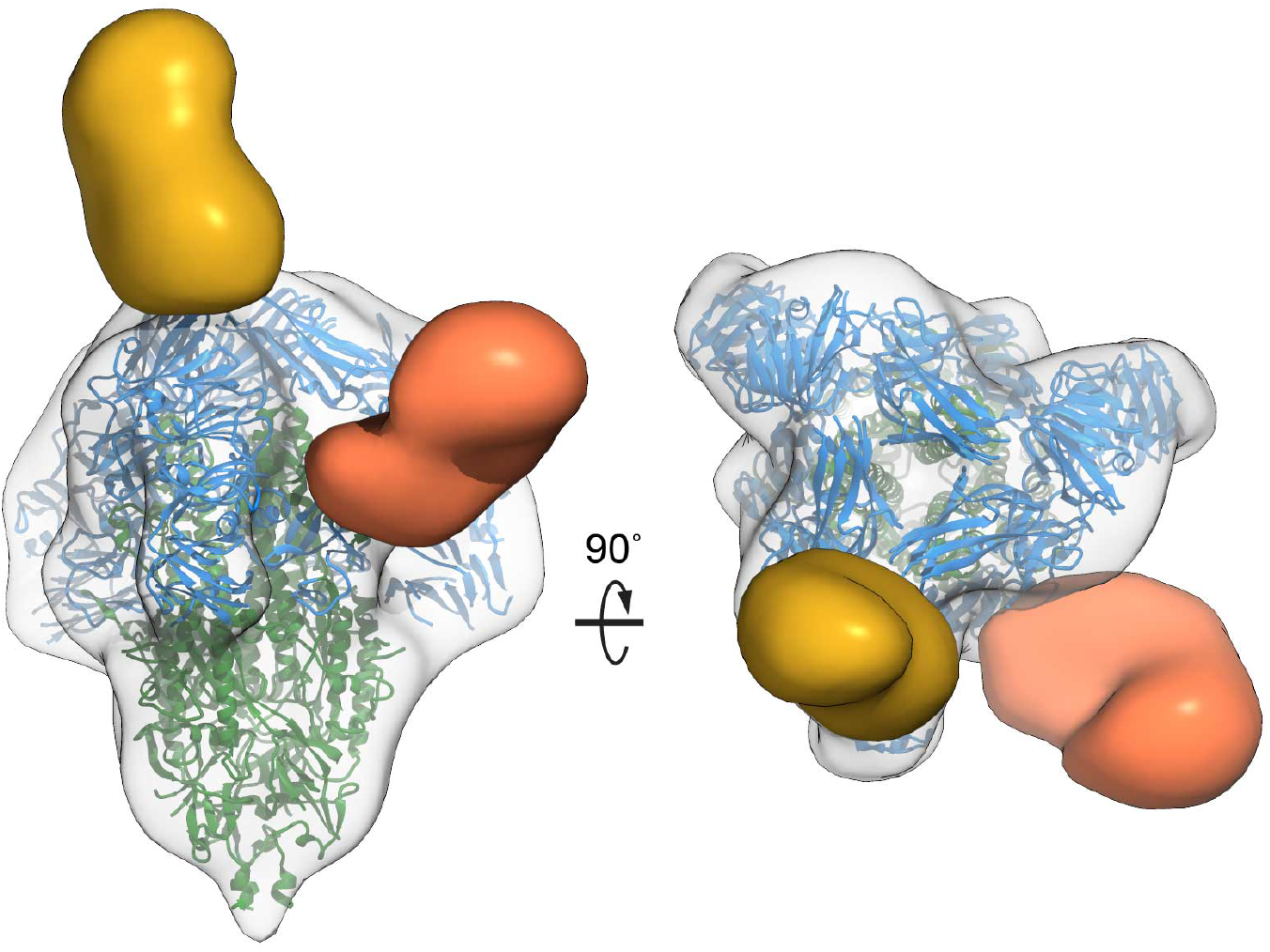
Polyclonal antibody recognition of PEDV spike epitopes. Antibody Fabs from three experimentally PEDV infected pigs recognize two distinct epitopes. The first epitope (yellow) is at the spike apex and is composed primarily of S1 domain A. The second epitope (orange) is on the side of the spike protein and is composed of S1 domains C and D and corresponds to a known neutralizing epitope (Okda et al., 2017; Sun et al., 2007).

## Discussion

These structures of the PEDV spike visualized by electron microscopy present new opportunities for structural comparisons against the collection of available coronavirus spike structures as well as indicating unique features such as glycan and fatty acid ligands mediating domain interfaces. We also determined the dominant antibody epitopes on the PEDV spike, one of which overlaps with a known neutralizing antibody epitope.

Similar to NL63 spike, the PEDV spike domain B are all in downwards conformations. This is in contrast to the betacoronavirus spike domain B of SARS- and MERS-CoV which adopt upwards conformations to recognize host protein receptors (Kirchdoerfer et al., 2018; Song et al., 2018). It has previously been proposed that the PEDV spike may use porcine APN as a host protein receptor for viral entry. However, recent evidence suggests APN is not necessary for PEDV entry into host cells (Li et al., 2017b). The observed poor accessibility of the PEDV spike domain B for receptor recognition supports the notion that this domain may recognize host receptors more poorly than the domain B of spikes which transition to upwards conformations more readily.

Spike conformational changes like those induced by host receptor binding (Walls et al., 2019), initiate the transition of coronavirus spikes from a pre-fusion to a postfusion state during viral entry. Key to these conformational transitions is the stability between the S1 and S2 regions of the viral spike as S1 shedding has been suggested as a prerequisite for this transition (Walls et al., 2017; Yuan et al., 2017). Our observation of glycan and fatty acid ligands in S1-S2 interfaces suggests a role for these ligands in modulating these interfaces and we hypothesize that these glycan and fatty acid interactions serve to stabilize the viral spike. These non-protein components add new considerations to protein engineering efforts to stabilize the prefusion conformations of alphacoronavirus spikes to take into account the effect of these non-protein ligands and their contribution to S1-S2 interactions.

In considering vaccine immunogens, it is also important to consider those epitopes which are dominantly recognized by the host immune response. The analysis of sera from pigs experimentally infected with PEDV revealed the same two epitopes in all three experimentally infected pigs studied, demonstrating a remarkably convergent immune response to infection. The epitope at the junction of domains C and D is a known neutralizing epitope (Okda et al., 2017; Sun et al., 2007). Moreover, previous work on this epitope demonstrated that immunizing with just this region of the PEDV spike was sufficient to produce sera with significant protection against viral infection (Sun et al., 2007). In addition, the paucity of diverse epitopes seen in the PEDV infected pig sera samples suggests significant opportunity for the specific presentation of spike protein regions as vaccine immunogens to explore and exploit novel epitopes not dominant in natural infection particularly against the S2 region of the protein.

These analyses of the PEDV spike reveal the ectodomain protein structure, non-protein ligands contributing to protein stability and dominant spike antibody epitopes arising during PEDV infection. Each of these observations will find future utility in not only understanding PEDV entry and immune responses, but also aid in the design of vaccine immunogens across the alphacoronavirus family.

## Acknowledgments

We thank Charles A. Bowman and Jean-Christophe Ducom for computational support, Hannah L. Turner and Bill Anderson for microscopy support and Lauren G. Holden for reviewing this manuscript. This work was supported by the National Institutes of Health, National Institute of Allergy and Infectious Disease AI123498 to R.N.K. and AI127521 to A.B.W.

## Author Contributions

R.N.K, L.M.S. and A.B.W. designed experiments. R.N.K and O.M. carried out sample preparation for structural analysis, data collection and processing and coordinate refinement. R.N.K, L.M.S, S.B. and A.B.W. analyzed data. M.B. and K.Y. provided porcine polyclonal antibody samples targeting the PEDV spike. All authors wrote and approved the final manuscript.

## STAR Methods

### LEAD CONTACT AND MATERIALS AVAILABILITY

Further information and requests for resources and reagents should be directed to and will be fulfilled by the Lead Contact, Andrew B. Ward. Plasmids for the expression of PEDV spike in mammalian and insect cells are available from the lead contact. There are restrictions to the availability of porcine sera due to limited availability.

### EXPERIMENTAL MODEL AND SUBJECT DETAILS

#### E. coli DH1OBac

*Escherchia coli* for the formation of recombinant bacmids, DH10Bac (Genotype: F–mcrA Δ(mrrhsdRMS-mcrBC) ϕ80lacZΔM15 ΔlacX74 recA1 endA1 araD139 Δ(ara,leu)7697 galU galK λ–rpsL nupG/bMON14272/pMON7124) carries the recombinant baculovirus genome, *Autographica californica,* on a bacterial artificial chromosome. These cells are propagated at 37°C in LB broth or on LB agar supplemented with 50 μg/mL kanamycin, 10 μg/mL tetracycline.

#### Sf9 Cell line

Sf9 cells (Expression Systems, Cat. #94-001S) are derived from Spodoptera frugiperda and are used for the production of recombinant baculoviruses. Cells are propagated in ESF-921 media (Expression Systems, Cat. #96-001-01) at 27°C in suspension in unbaffled Erlenmeyer flasks or in adherent cultures. Cells are maintained at a density between 0.8-4.0×10^6^ cells/mL. The cells are female.

#### 293F Cell line

293F cells are a 293 human embryonic kidney cell line adapted for suspension culture in FreeStyle 293 Expression Medium (Thermo Fisher, Cat #12338026). Cells are maintained at 37°C in 8% CO2 in suspension. Cells are grown in smaller cultures less than 150 mL in unbaffled Erlenmeyer flasks or in larger cultures in baffled Erlenmeyer flasks where cells are maintained between 0.3-2.0×10^6^ cells/mL. Cells are female.

#### 293S Cell line

293S cells are a 293 human embryonic kidney Gnt knockout cell line adapted for suspension culture. Cells are maintained in FreeStyle 293 Expression Medium. Cells are maintained at 37°C in 8% CO_2_ in suspension. Cells are grown in smaller cultures less than 150 mL in unbaffled Erlenmeyer flasks or in larger cultures in baffled Erlenmeyer flasks where cells are maintained between 0.3-2.0×10^6^ cells/mL. Cells are female.

#### Sus scrofa

The experimental design was approved by the University’s Institutional Animal Care and User Committee (protocol log #6-13-7593-S). Three-week old, cross-bred, weaned pigs of mixed sex (female and barrow) were purchased from a single commercial farrow-to-wean herd without any known prior exposure to PEDV, porcine reproductive and respiratory syndrome virus (PRRSV) and transmissible gastroenteritis virus (TGEV). Seronegative status for all these viruses was also confirmed by laboratory testing before receiving. The pigs were kept in the Livestock Infectious Disease Isolation Facility (BSL2 compliance) at Iowa State University (Ames, Iowa) and randomly assigned to challenged or sham-control group each of which was separated by room and ventilation system in the same facility. Pigs in each room were confined by pens on a solid floor that was rinsed daily, fed a balanced diet *ad libitum* based on weight, and given free access to water.

## METHOD DETAILS

### Plasmid preparation

The protein sequence of Porcine epidemic diarrhea virus (PEDV) USA/Colorado/2013 spike ectodomain 1-1322 with a C-terminal foldon, TEV site, His-tag and Strep-tag was codon optimized for mammalian expression (Genscript) and inserted into pFastBac1 (Thermo Fisher, Cat. #10360014) using EcoRI and XbaI restriction sites (Genscript). The insert was further cloned into pcDNA3.4 using the HiFi DNA Assembly Kit (New England Biolabs) and PCR products (Phusion polymerase, Thermo Fisher, Cat. #F530L) for spike (PEDV fwd and rvs) and for pcDNA3.4 (pcDNA34 fwd and rvs).

The Asn264Asp in spike-pcDNA3.4 substitution and 34-230 deletion spike (based on genbank sequence AMK69964) in spike-pFastBac1 mutants were made by Phusion PCR of the parent templates using N264D fwd and rvs or PEDVS0033 rvs and PEDVS0231 fwd primers. The resulting PCR products were purified with NucleoSpin PCR cleanup kit (Machery Nagel, Cat. #740609.50) to which was added 10X PNK buffer, 1 μL T4 polynucleotide kinase and 1 μL DpnI (New England Biolabs, Cats. #M0201S and R0176S) and incubated at 37°C for 1 hour. 3μL of this reaction, 1μL of 10X T4 ligase buffer, 1μL of T4 ligase (New England Biolabs, Cat. #M0202S) and 5μL of water were incubated at room temperature for 1 hour. The ligated DNA was then transformed into DH5α *E. coli* (Thermo Fisher, Cat. #18258012) and selected on LB broth agar with 100μg/mL ampicillin.

### Production of recombinant baculoviruses

Protein expression constructs in pFastBac1 were transformed into DH10Bac *E. coli* (Thermo Fisher, Cat. # 10361012) and selected on Luria broth agar with 50μg/mL kanamycin, 15μg/mL gentamycin, 0.5 mM isopropylthiogalactopyranoside (IPTG) and 20 μg/mL X-gal at 37°C. White colonies, indicating recombinants, were grown in LB broth and the recombinant bacmid DNA was prepared using the NucleoBond Xtra Bac Kit (Machery Nagel, Cat. #740436.25). 1μg of recombinant bacmid DNA was combined in 1 mL of ESF-921 media (Expression Systems, Cat. #96-001-01) with 20 μL of Cellfectin II (Thermo Fisher, Cat. #10362100) and added to 1×10^6^ adherent Sf9 cells. Cells were incubated for five days. Cell supernatants were clarified by centrifugation at 1500×g for 5 minutes. 0.5 mL of transfected cell supernatant was used to amplify the baculoviruses on 5×10^6^ adherent Sf9 cells for 5 days. First virus passage supernatants were clarified by centrifugation. 1 mL of first passage supernatant was used to further amplify the baculovirus on 50 ×10^6^ suspension Sf9 cells and incubated for 5 days.

### Recombinant protein production and purification

Baculovirus expressed proteins were produced by infecting 1 L of Sf9 at 2×10^6^ cells with 10 mL of second amplification baculovirus and incubating for four days. Infected cell supernatants were harvested by centrifugation at 1500×g to remove cells followed by 12,000 ×g to remove cell debris. Supernatants were filtered through a 0.45 μm filter and then applied to a column of Streptactin agarose (Qiagen, Cat. #30004). The column was washed with 20 column volumes of Tris-buffered saline and then the protein was eluted with same buffer with 2 mM desthiobiotin. The eluted protein was purified by Superose6 Increase 10/30 (GE Life Sciences, Cat. #29091596) gel filtration.

Mammalian expressions were done by transient transfection. For both 293F and 293S cells, 1 L of cells at 1×10^6^ cells were transfected with complexes composed of 250 μg of plasmid DNA and 750 μg of polyethylenimine (Polysciences, Cat. # 24765-1). For the expression involving kifunensine, three hours after transfection kifunensine (Cayman Chemicals, Cat. #10009437) was added to a final concentration of 5 μM. Expressions were allowed to proceed for five days. Cell supernatants were collected by centrifugation at 1,500 × g to remove cells followed by 12,000 × g to remove cell debris. Supernatants were filtered through a 0.45 μm filter and then applied to a column of Streptactin superflow. The column was washed with 20 column volumes of Tris-buffered saline and then the protein was eluted with same buffer with 2 mM desthiobiotin. The eluted protein was purified by Superose6 Increase 10/300 gel filtration.

Proteins were concentrated for structural studies using ultrafiltration (Sigma-Millipore, AmiconUltra 4 mL, 30K MWCO, Cat. #UFC903024).

### PEDV infection and sera collection

After 3-day acclimation, pigs in the challenged group were inoculated with a prototype US PEDV isolate US/Iowa/18984/ 2013 (GenBank accession #KF804028) (Hoang et al., 2013), 10^3^ plaque-forming units/pig, via oro-gastric gavage as previously described (Madson et al., 2014). Pigs in the control group received virus-free cell culture media. Some of these pigs were inoculated again with the same strain in the identical manner at day 56 post-inoculation (PI). The positive swine sera used in the study were collected from pigs on day 76 PI (i.e., 20 days after the 2nd inoculation). The negative swine serum was from one of age-matched control pigs. The positive serum had an antibody titer of 8 to 32 against PEDV as measured by serum-virus neutralization test.

### Preparation of polyclonal Fab fragments

IgG was isolated from porcine sera by passing the sera over a MabSelect 5 mL HiTrap Column (GE Life Sciences), washing with two column volumes of tris-buffered saline and elution with 0.1 M glycine, pH 2.5. Acid elutions were immediately neutralized with 1/10^th^ volume of 1M tris base and then buffer exchanged 1:10 twice into tris-buffered saline. IgG was digested to Fab and Fc using papain. Digestions were done in tris-buffered saline (25mM TrisCl pH 7.4, 150 mM NaCl) with 5 mM dithiothreitol using 0.25% (w/w) papain for 2 hours at room temperature. The digest was quenched by diluting 200 mM iodoacetamide (Sigma) 1/20 into the reaction. Undigested IgG was removed from the quenched reactions by passing the reactions over a Superdex200 Increase 10/300 column (GE Life Sciences, Cat. #28990944) in tris-buffered saline.

For polyclonal epitope analysis, 30 μg of purified Sf9 expressed full-length PEDV spike ectodomain was combined with 1.5 mg of Fab/Fc fractions from gel filtration. The proteins were incubated overnight at room temperature. Fab-spike complexes were isolated by gel filtration using a Superdex200 Increase 10/300 column in tris-buffered saline monitoring the elutions at 215 nm and collecting the spike peaks. Collected fractions were concentrated by ultrafiltration.

### Negative-stain electron microscopy

PEDV spikes and spike-Fab complexes for negative-stain electron microscopy were diluted to 30-50 μg/mL using tris-buffered saline. 3 μL of protein sample was spotted onto glow-discharged carbon-coated copper grids and stained with uranyl formate. Imaging of grids was performed using a T12 Tecnai Spirit operating at 120 kV and Tietz 4K camera and Leginon (Suloway et al., 2005) data collection software. Images were assesed using EMHP (Berndsen et al., 2017) and processed using RELION-3.0 (Zivanov et al., 2018) and CryoSparc 2.0 (Punjani et al., 2017). Briefly for spike samples, particles were extracted in RELION binning by 2 and then reconstructed in CryoSparc. For spike-Fab complexes, particles were extracted binning by 2 and reconstructed in C1 in RELION before extensive 3D classification with local alignment followed by C1 3D refinement of unique classes.

### Cryo-electron microscopy

PEDV spike ectodomain for cryo-EM was used at a final concentration of 1.3 mg/mL with 0.01% amphipol A8-35 (Anatrace). The protein was spotted onto UltraAuFoil 1.2/1.3 gold grids (Quantifoil) plasma cleaned using an Ar/O_2_ gas mix. Grids were blotted for 3.5 s using a Vitrobot IV (Thermo Fisher) and then plunge frozen into liquid ethane at −185°C. Grids were imaged using a Talos Arctica (Thermo Fisher) at 200 kV with a K2 Summit camera (Gatan) and Leginon data collection software. Micrograph movies were aligned using MotionCor2 (UCSF) (Zheng et al., 2017), CTF corrections were done with Gctf (Zhang, 2016) and images were assessed with EMHP (Berndsen et al., 2017). Further data processing was performed in RELION-3.0 (Zivanov et al., 2018). Particles were picked, extracted, 2D classified and 3D refined. Attempts at C1 refinements and 3D sorting all produced reconstructions resembling C3 symmetry and so C3 symmetry was used in the final reconstruction.

The model of NL63 spike ectodomain (5SZS.pdb) (Walls et al., 2016b) was used as a starting model. The model was aligned an mutated to the PEDV spike sequence and rebuild using Coot (Emsley et al., 2010). Coordinate models were further refined using real-space refinement in Phenix (Adams et al., 2010) and RosettaRelax (Wang et al., 2016). Model validation was done using Molprobity (Williams et al., 2018) and EMRinger (Barad et al., 2015).

### Mass spectrometry

The PEDV spike sample expressed in Sf9 cells was analyzed on Zorbax SB-C18 column (Agilent) with water with 0.1% formic acid and acetonitrile with 0.1% formic acid as mobile phases. The mass spectrometry data was acquired in positive mode on Qtof Impact II (Bruker) and analyzed using Compass Data analysis 4.3 (Bruker).

### QUANTIFICATION AND STATISTICAL ANALYSIS

Statistical models inherent to RELION-3.0 (Zivanov et al., 2018) and CryoSparc 2.0 (Punjani et al., 2017) were used to create 3D classifications and 3D refinements from spike particle images.

### DATA AND CODE AVAILABILITY

All map and coordinate data presented in the manuscript has been deposited in Protein Data Bank (www.rcsb.org) with accession 6VV5 or in the Electron Microscopy Data Bank (www.ebi.ac.uk/pdbe/emdb/) with accessions 21391, 21392, 21393, 21394, 21395, 21396, 21397, 21398, 21399, 21400, 21401, 21402, 21403, 21404.

## KEY RESOURCES TABLE

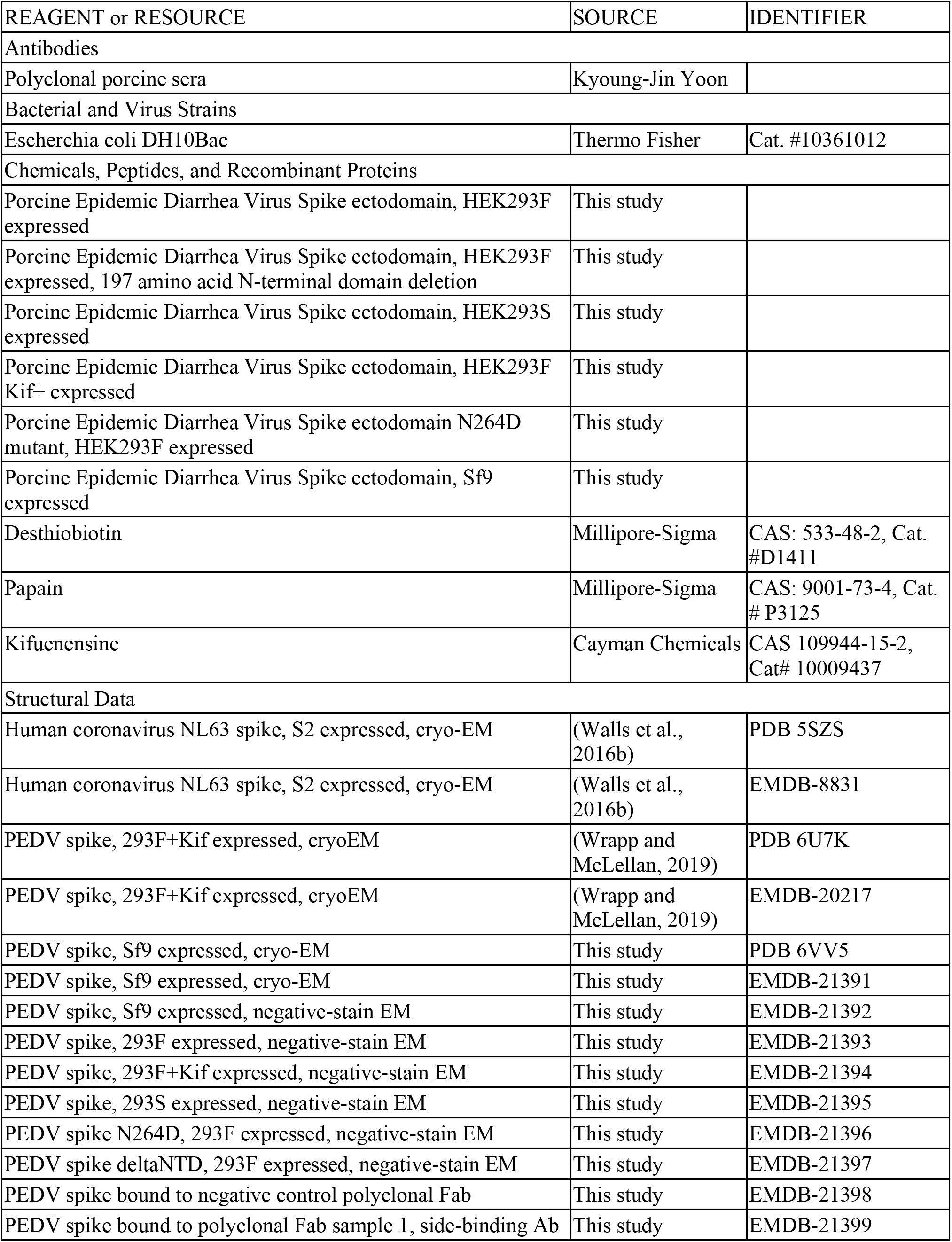
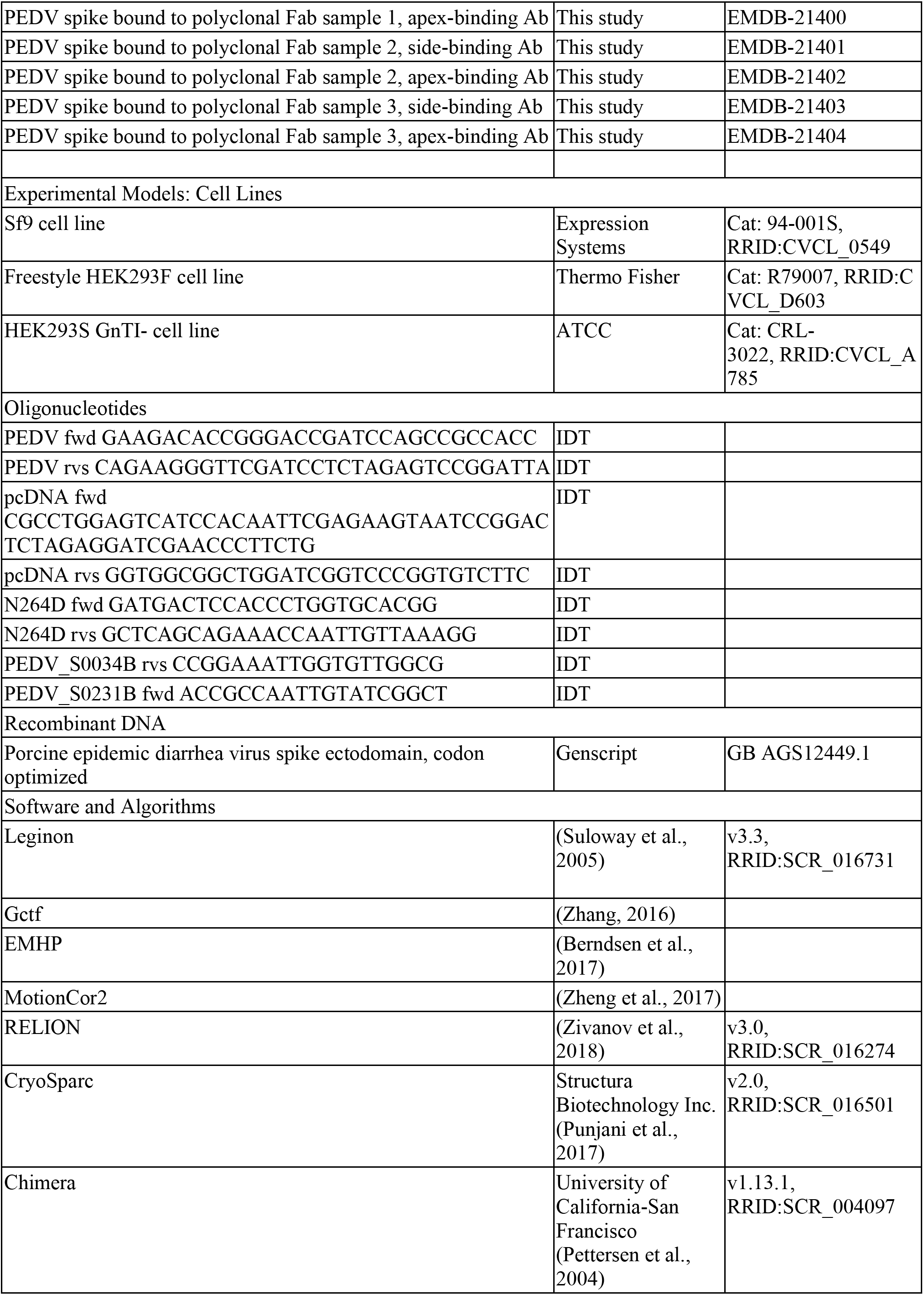
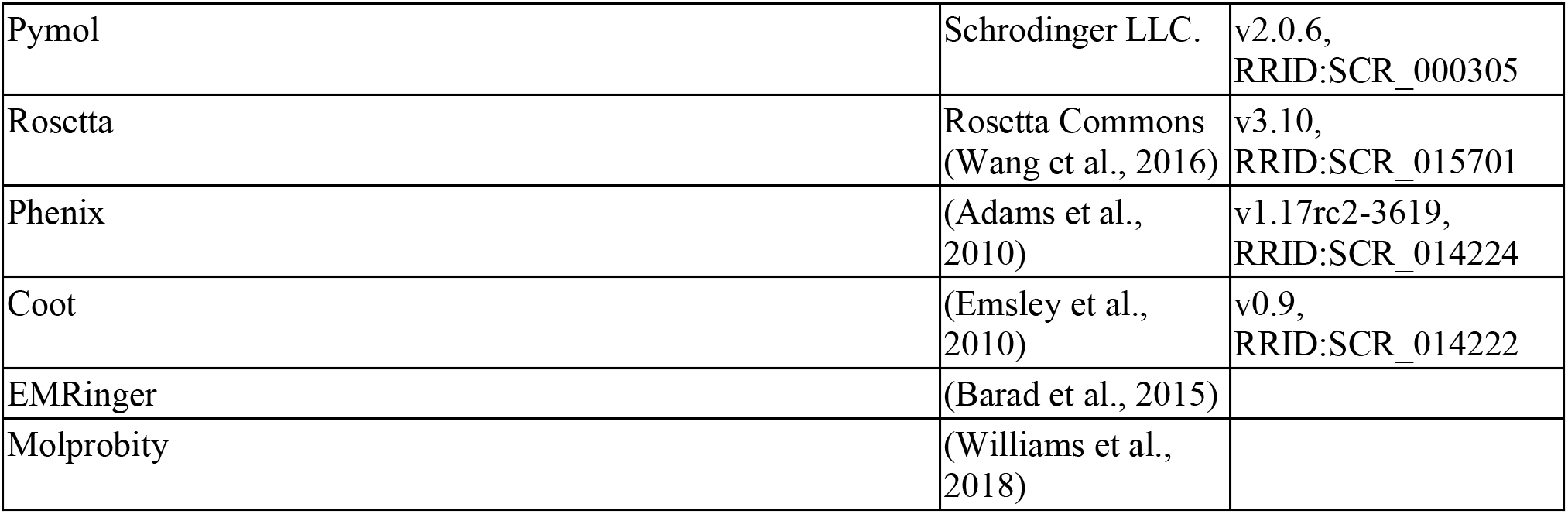

## Supplemental Information

**Supplemental Table 1:** Data collection and refinement of PEDV spike by cryo-electron microscopy.

|  |  |
| --- | --- |
| EMDB | 21391 |
| **Data and reconstruction statistics** |  |
| Microscope | Talos Actica |
| Voltage (kV) | 200 |
| Detector | Gatan K2 Summit |
| Dose rate (e<sup>-</sup>/pix/sec) | 5.67 |
| Exposure (s) | 12.8 |
| Dose (e<sup>-</sup>/Å<sup>2</sup>) | 54.9 |
| Frames | 64 |
| Defocus Range (μm) | 0.6-1.4 |
| Initial Particles | 19,436 |
| Final Particles | 19,436 |
| B-factor (Å<sup>2</sup>) | -70 |
| Resolution (Å) | 3.5 |
| **Coordinate model refinement** |  |
| PDB | 6VV5 |
| Residues | 3,396 |
| RMSD Bonds (Å) | 0.017 |
| RMSD Angles (°) | 1.579 |
| Ramachandran |  |
| Favored (%) | 98.3 |
| Allowed (%) | 1.6 |
| Outliers (%) | 0.1 |
| Rotamer Outliers (%) | 1.3 |
| Clash score | 4.8 |
| Molprobity score | 1.32 |
| EM Ringer Score | 3.3 |

**Supplemental Table 2:** Data collection for PEDV spikes by negative-stain electron microscopy. All data was collected on a Technai G2 Spirit operating at 120 kV with a Teitz 4K × 4K camera and collecting a total dose of 25 e^-^/Å^2^.

|  |  |  |  |  |  |  |
| --- | --- | --- | --- | --- | --- | --- |
| Sample | PEDV S 1-1322 | PEDV S 1-1322 | PEDV S 1-1322 | PEDV S 1-1322, N264D | PEDV S 1-1322 | PEDV S 1-33, 231-1322 |
| Expression System | 293F | 293F+Kif | 293S | 293F | Sf9 | Sf9 |
| EMDB | 21393 | 21394 | 21395 | 21396 | 21392 | 21397 |
| Pixel size | 2.05 | 2.05 | 2.05 | 2.05 | 2.05 | 4.1 |
| Particles | 29910 | 39819 | 11268 | 10698 | 12896 | 18070 |
| Resolution | 12 | 10 | 13 | 12 | 13 | 23 |

**Supplemental Table 3:** Data collection for PEDV spikes and porcine F_ab_ by negative-stain electron microscopy. All data was collected on a Technai G2 Spirit operating at 120 kV with a Teitz 4K × 4K camera and collecting a total dose of 25 e^-^/Å^2^.

| Sample | Negative control | Sample 1, side-binding | Sample 1, apex-binding | Sample 2, side-binding | Sample 2, apex-binding | Sample 3, side-binding | Sample 3, apex-binding |
| --- | --- | --- | --- | --- | --- | --- | --- |
| Expression System | Sf9, Sus Scrofa | Sf9, Sus Scrofa | Sf9, Sus Scrofa | Sf9, Sus Scrofa | Sf9, Sus Scrofa | Sf9, Sus Scrofa | Sf9, Sus Scrofa |
| EMDB | 21398 | 21399 | 21400 | 21401 | 21402 | 21403 | 21404 |
| Pixel size | 4.1 | 4.1 | 4.1 | 4.1 | 4.1 | 4.1 | 4.1 |
| Particles | 44484 | 4552 | 753 | 6504 | 5903 | 36978 | 15570 |
| Resolution | 15 | 22 | 28 | 23 | 23 | 15 | 17 |

**Supplemental Figure 1:**
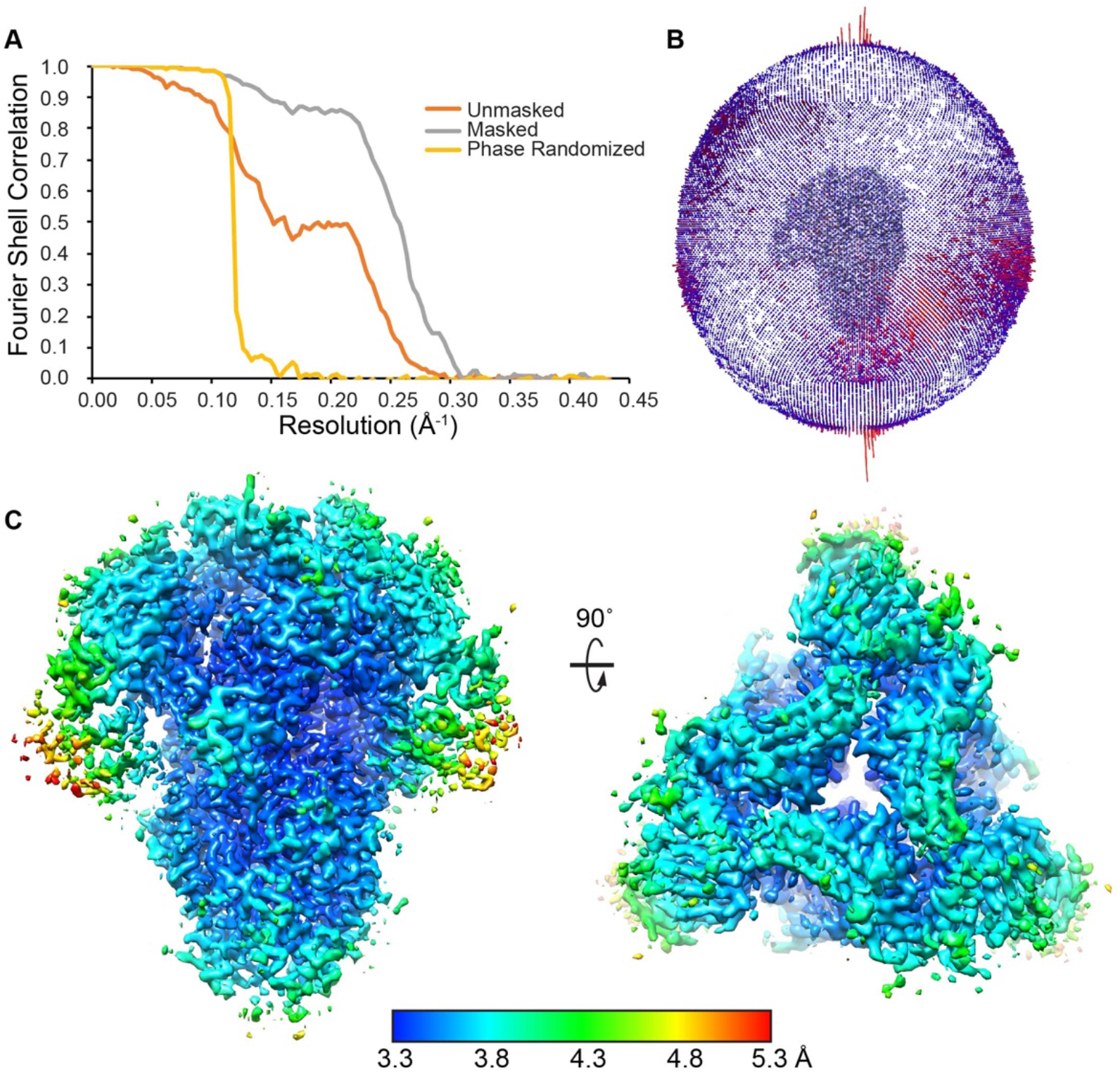
Validation of cryo-electron microscopy reconstruction. A) Fourier shell correlation curves for unmasked, masked and phase randomized correlations. B) Plot of angular distribution. C) Local resolution of the reconstruction calculated with RELION-3.0 (Zivanov et al., 2018).

**Supplemental Figure 2:**
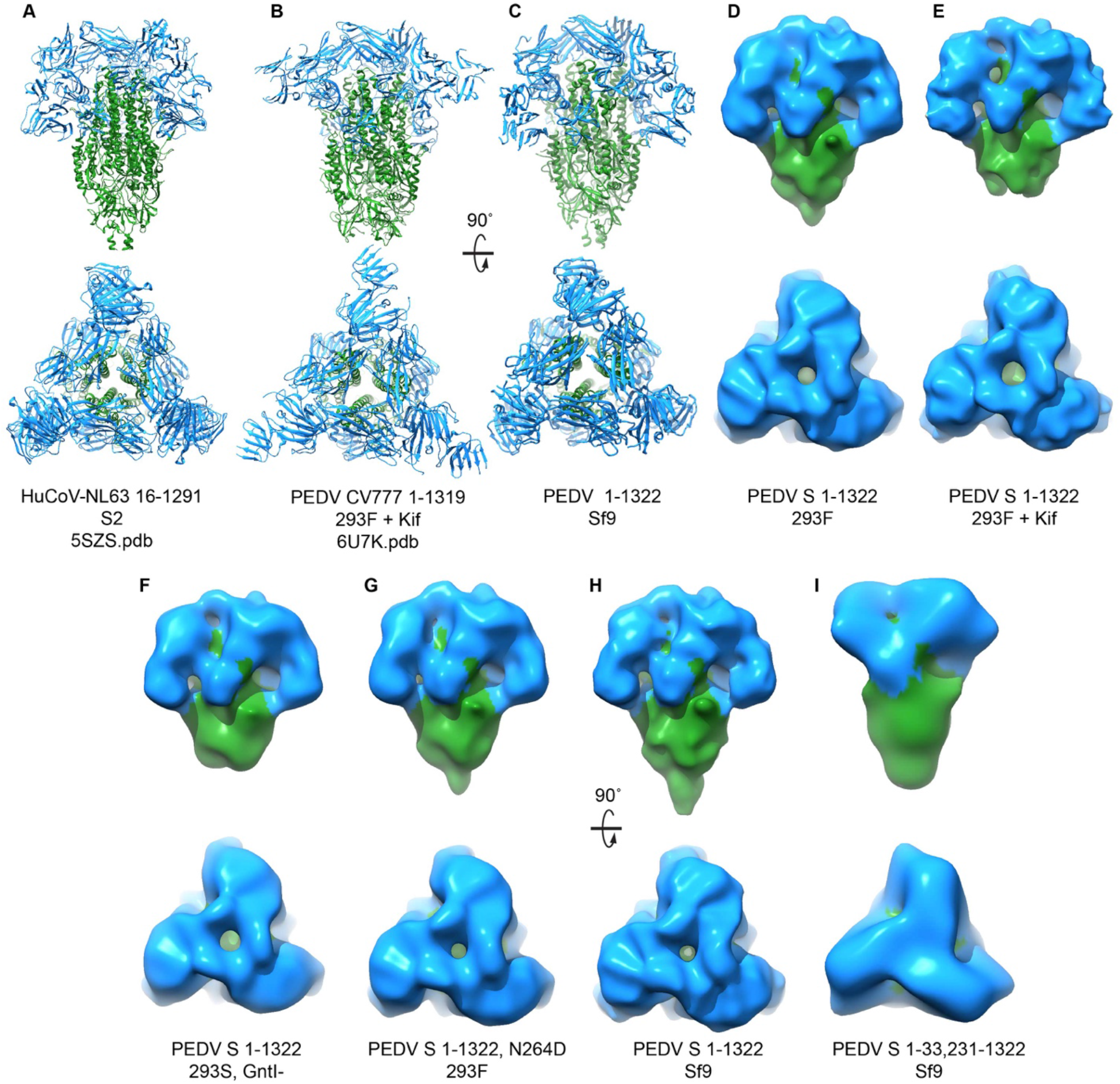
Conformations of Alphacoronavirus spike variants. A) The HuCoV-NL63 amino acids 16-1291 spike cryoEM structure expressed from Drosophila melanogaster S2 cells. B) The previously published PEDV spike cryoEM structure, strain CV777, amino acids 1-1319 expressed in human 293F cells with kifunensine. C) The PEDV spike cryoEM structure, amino acids 1-1322 expressed in Sf9 cells. D) PEDV spike was expressed in 293F cells, adding complex glycans. E) PEDV spike was expressed in 293F cells in the presence of kifunensine adding high-mannose glycans. F) PEDV spike was expressed in 293S GntI-cells adding high-mannose glycans. G) PEDV spike carrying a knockout of the glycan at Asn264 was expressed in 293F cells. H) PEDV spike was expressed in Sf9 insect cells, adding short high-mannose glycans. I) PEDV spike was truncated to remove the first N-terminal domain, domain 0 (amino acids 34-230) and expressed in Sf9 insect cells. With the exception of (B), all PEDV spike structures are of strain USA/Colorado/2013.

**Supplemental Figure 3:**
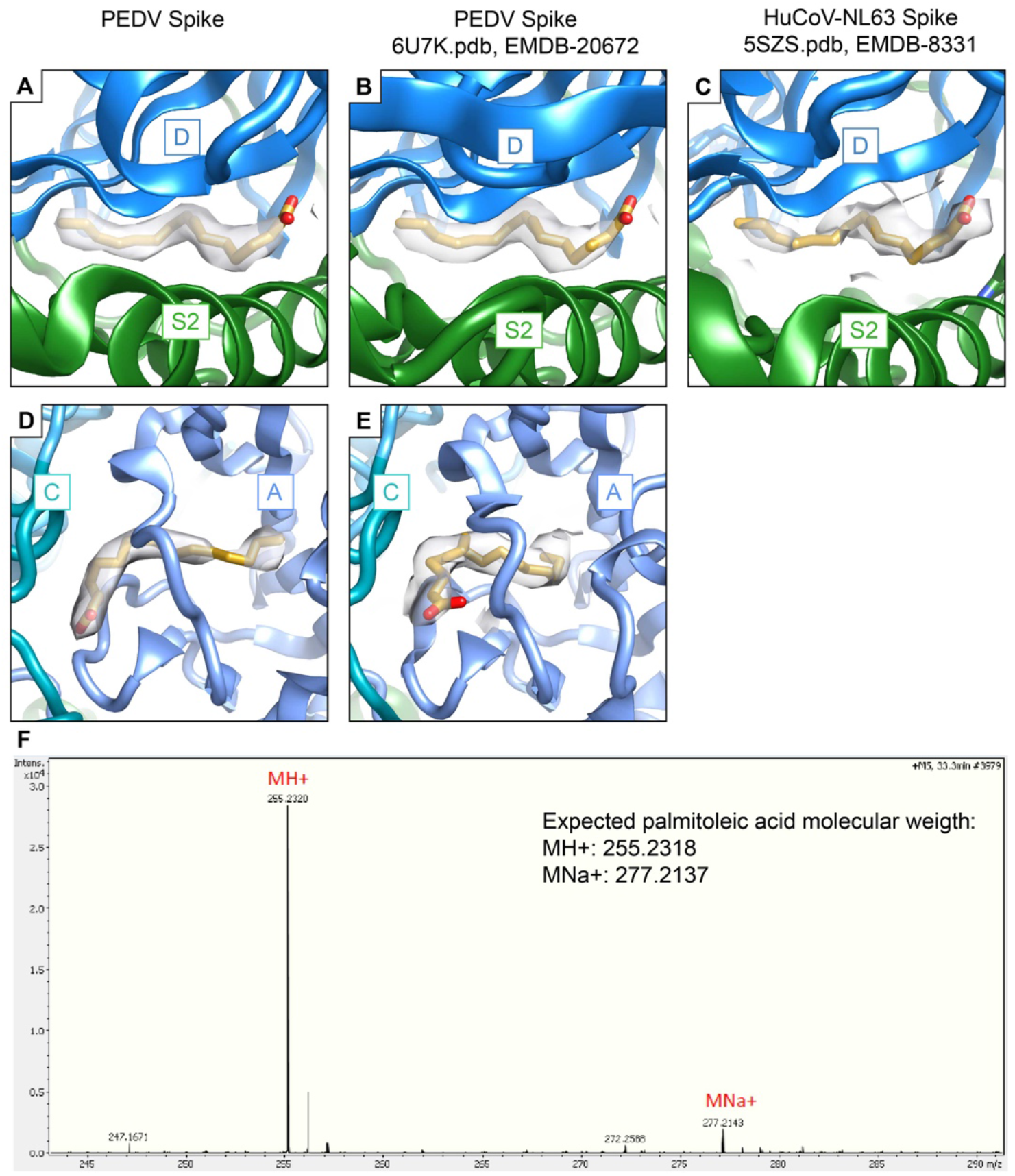
Alphacoronavirus spike proteins bind to palmitoleic acid. Comparison of the palmitoleic acid binding site in density maps for A) the PEDV spike structure determined here, B) the previously determined PEDV spike structure (Wrapp and McLellan, 2019) and C) the previously determined HuCoV-NL63 spike structure (Walls et al., 2016b). The coordinates of palmitoleic acid built in the PEDV spike structure presented here were superposed onto the other spike structures based on neighboring protein regions. Comparison of density in the second palmitoleic acid binding sites located within PEDV S1 domain A in D) the spike structure presented here as well as E) the previously determined PEDV spike structure (Wrapp and McLellan, 2019). Note that the palmitoleic acid density appears to adopt an alternate conformation in the previously determined PEDV spike structure. The coordinates of palmitoleic acid were real space refined to reflect this altered conformation. The altered fatty acid conformation may be due to differences in the conformation of this region of the protein (see main text).

**Supplemental Figure 4:**
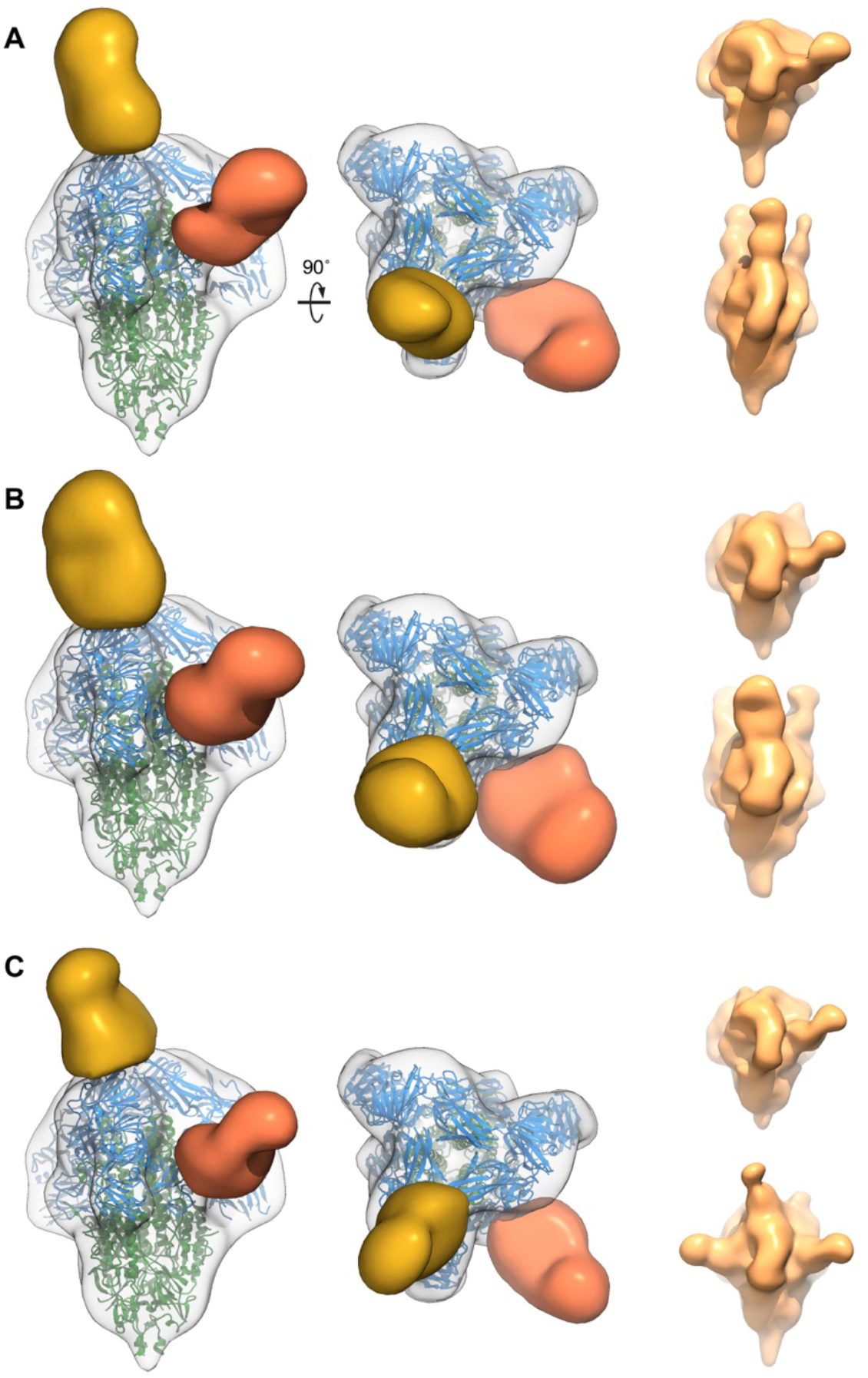
Polyclonal antibody analysis of pigs experimentally infected with PEDV. The polyclonal antibody sera from pigs experimentally infected with PEDV was bound to PEDV spikes and analyzed by negative-stain electron microscopy. Segmented maps of antibody Fabs from three individual pigs (A, B and C) confirm the recognition of the same two antibody epitopes on PEDV spikes. For each sera sample, representative non-segmented maps are shown at right.

